# Conformational dynamics and activation of membrane-associated human Group IVA cytosolic phospholipase A_2_ (cPLA_2_)

**DOI:** 10.1101/2025.03.22.644760

**Authors:** Mac Kevin E. Braza, Edward A. Dennis, Rommie E. Amaro

## Abstract

Cytosolic phospholipase A_2_ (cPLA_2_) associates with membranes where it hydrolyzes phospholipids containing arachidonic acid to initiate an inflammatory cascade. All-atom molecular dynamics simulations were employed to understand the activation process when cPLA_2_ associates with the endoplasmic reticulum (ER) membrane of macrophages where it acts. We found that membrane association causes the lid region of cPLA_2_ to undergo a closed-to-open state transition that is accompanied by the sideways movement of loop 495-540, allowing the exposure of a cluster of lysine residues (K488, K541, K543, and K544), which binds the allosteric activator PIP_2_ in the membrane. The active site of the open form of cPLA_2_, containing the catalytic dyad residues S228 and D549, exhibited a three-fold larger cavity than the closed form of cPLA_2_ in aqueous solution. These findings provide mechanistic insight as to how cPLA_2_ ER membrane association promotes major transitions between conformational states critical to allosteric activation and enzymatic phospholipid hydrolysis.

## Main text

The interaction between cytosolic proteins and bilayer membranes of intracellular organelles is central to biological function and signaling. Integrating cellular, biochemical, and structural information is of the utmost importance in elucidating protein-membrane interactions at the atomistic level.^1,2^ In the current study, we investigated how the human cytosolic phospholipase A_2_ (cPLA_2_),^3^ a critical enzyme in the inflammatory cascade,^2^ associates with intracellular endoplasmic reticulum (ER) membranes, where it undergoes a critical state transition. The presence of known allosteric regulators such as membrane association, Ca^2+^, and the phosphatidylinositol (4,5)-bisphosphate (PIP_2_) were also elucidated.

Phospholipase A_2_ (PLA_2_) constitutes a superfamily of enzymes that hydrolyze the fatty acyl group occupying the *sn*-2 position of membrane phospholipids.^3^ The Group IVA (GIVA) cPLA_2_ exhibits specificity for phospholipids containing arachidonic acid (AA) at the *sn*-2 position.^4^ This specificity can be explained at the molecular level, where several aromatic amino acid side chains form a precise binding site cavity for AA. AA consists of 20 carbons and four cis double bonds that exhibit π-π stacking with the aromatic amino acid side chains in the binding site.^4^ At the cellular level, cPLA_2_ is a water-soluble protein that associates with the surface of the phospholipid bilayer. Once cPLA_2_ pulls a single phospholipid substrate into its active site, it hydrolyzes and releases it, whereby downstream enzymes bind the AA and initiate the inflammatory cascade.^2^ To date, there is only one crystal structure of the Group IVA cPLA_2_.^5.^ It contains both a C2 and a catalytic domain, but a mechanistic understanding of the synergism between these two domains is lacking. Previous studies using both classical and steered molecular dynamics (MD) simulations revealed that the lid region of the membrane-associated cPLA_2_ exhibited high flexibility before substrate binding of 1-palmitoyl-2-arachidonoyl-*sn*-glycero-3-phosphocholine (PAPC or 16:0/20:4 PC).^4^

The Monod–Wyman–Changeux (MWC) model describes allosteric regulation, where distant ligand binding drives cooperative conformational changes.^6^ In the classical MWC allosteric protein nicotinic acetylcholine receptor (nAChR), Ca^2+^ modulates acetylcholine binding at distinct sites.^7,8^ Similarly, cPLA_2_ is allosterically regulated by Ca^2+^ (∼30 Å from the active site), with PIP_2_ and membranes also acting as allosteric regulators, as shown by the Dennis group.^1,4^

Hydrogen/Deuterium exchange-mass spectrometry (HDX-MS) data for proteins provides qualitative analysis by mapping the average exchange rates of peptides under varying conditions.^9–11^ Although HDX-MS can be used as an indirect probe of the protein allostery and dynamics, it can be limited. MD simulations help augment and extend the available HDX-MS, structural, dynamical, and biological data to provide an atomically detailed mechanism of protein dynamics. Furthermore, MD simulations can elucidate allosteric processes under different model types and conformational landscape shifting.^12,13^

In the present study, we carried out all-atom molecular dynamics simulations of the aqueous and membrane-associated cPLA_2_ to complement findings from *in vivo*, X-ray crystallography, lipidomics, and HDX-MS data. Furthermore, we analyzed the conformational dynamics induced by the three allosteric regulators, namely membrane association, Ca^2+^, and PIP_2_. The detailed computational methods section can be found in the Supplementary Information section. In brief, we analyzed three replicates of 500 ns explicit solvent classical MD trajectories for each system, gathering a combined 7.5 μs dataset (Table 1). In summary, the five systems we simulated were (1) aqueous cPLA_2_ with bound Ca^2+^, (2) membrane-associated cPLA_2_, (3) membrane-associated cPLA_2_ + Ca^2+^, (4) membrane-associated cPLA_2_ + PIP_2_; and (5) membrane-associated cPLA_2_ + Ca^2+^ + PIP_2_. Furthermore, we describe various conformational dynamics of the C2 and catalytic domain, lid region opening, and loop 495-540.

**Table 1.** cPLA_2_ -membrane systems employed in the present study.^a,b,c,d,e^

| System number | Ca<sup>2+</sup> | 1% PIP<sub>2</sub> | Number of lipids |  | Total no. of atoms | Simulation length |
| --- | --- | --- | --- | --- | --- | --- |
|  |  |  | Upper leaflet | Lower leaflet |  |  |
| 1 | + | - | - | - | 424,425 | 3 replicates x 500 ns |
| 2 | - | - | 400 | 400 | 495,230 | 3 replicates x 500 ns |
| 3 | + | - | 400 | 400 | 495,366 | 3 replicates x 500 ns |
| 4 | - | + | 400 | 400 | 495,388 | 3 replicates x 500 ns |
| 5 | + | + | 400 | 400 | 495,110 | 3 replicates x 500 ns |

<sup>a</sup>All systems were simulated using the CHARMM36 force field and built using the CHARMM-GUI Membrane Builder.
<sup>b</sup>Membrane compositions were derived from Andreyev et al.<sup>14</sup>
<sup>c</sup>Na<sup>+</sup> and Cl<sup>-</sup> ions at 150 mM were added.
<sup>d</sup>cPLA<sub>2</sub> Histidine protonation states were set based on the PROPKA output.
<sup>e</sup>cPLA<sub>2</sub> orientation to the membrane was consistent with the OPM and HDX-MS data.

We built the RAW264.7 macrophage ER membrane environment with its associated cPLA_2_ (Figure 1). The orientation of cPLA_2_ with the membrane is consistent with the predicted placement from the Orientations of Proteins in Membranes (OPM) database.^15^ Furthermore, this is consistent with all previous studies, especially the available cPLA_2_ HDX-MS data.^1,4,16–20^ The ER has been reported to be comprised of glycerophospholipids, sphingolipids, and sterols at varying concentrations. We used the subcellular lipidome data reported for the ER of RAW264.7 macrophages.^14^ Furthermore, it has been reported that PI is the most abundant 20:4-containing glycerophospholipid in RAW264.7 macrophages.^14,21^ In the most recent study of Murawska et al., they verified that the phospholipid 1-stearoyl-2-arachidonoyl phosphatidylinositol (SAPI or 18:0/20:4 PI) is the most specific substrate for cPLA_2_ in the RAW macrophages.^21^ The ER membrane contains 10% SAPI. The CHARMM-GUI membrane builder was used to embed cPLA_2_ in the membrane and built the extensive ER membrane.^22^ Although we recognized that the upper and lower leaflets of the ER membrane are probably asymmetric, for our purposes, symmetric membranes were employed in these studies. The exact composition of the lipids employed in the models is reported (Table S1 and Table S2).

**Figure 1.**
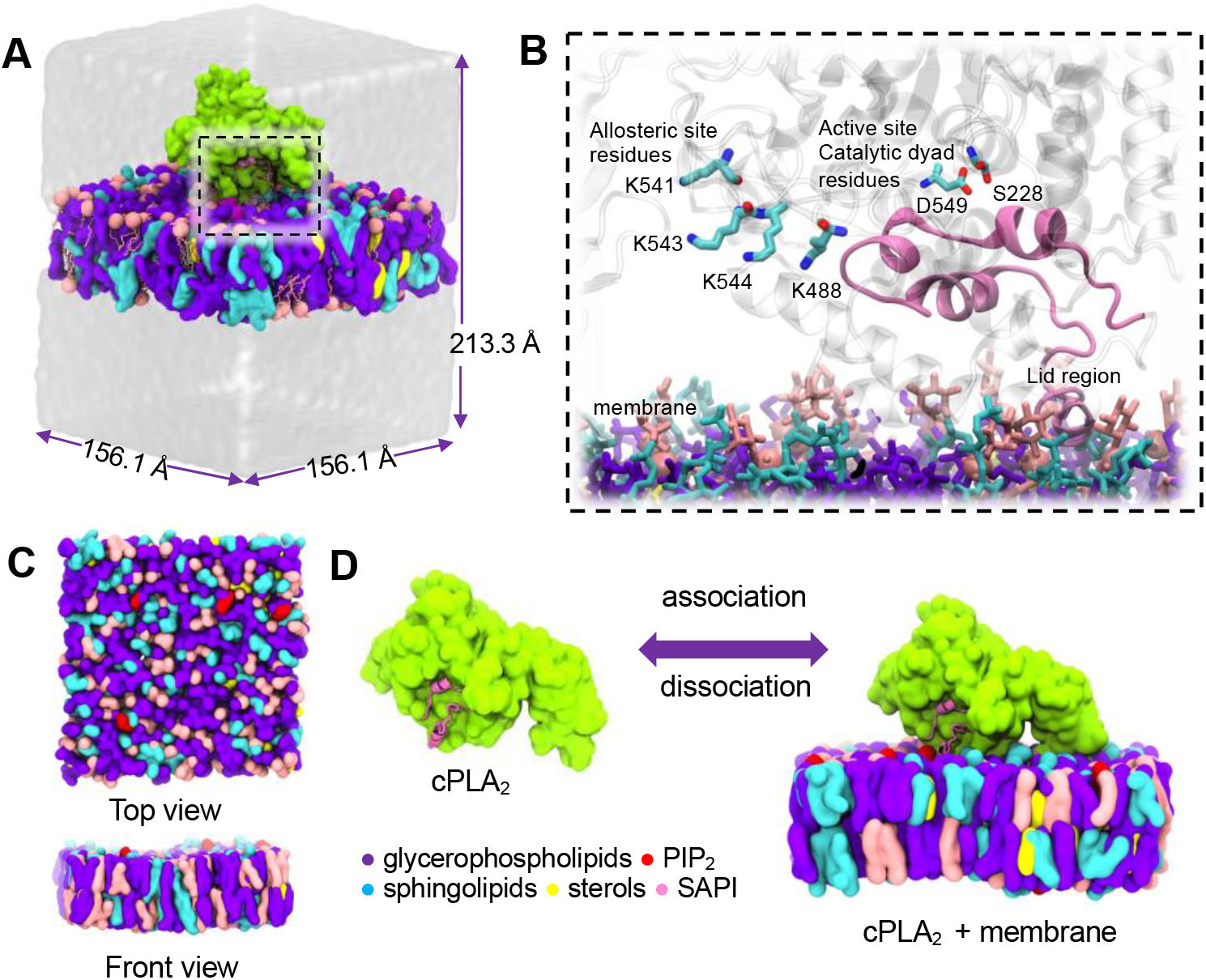
Human cytosolic phospholipase A_2_ (cPLA_2_) and RAW264.7 macrophage ER membrane interactions. (A) Cellular environment of cPLA_2._ (B) Phospholipid active site catalytic dyad residues (S228 and D549) and allosteric site residues K488, K541, K543, and K544. The lid region is also highlighted (magenta). (C) The membrane lipid bilayer model was assumed in this study with the exact lipid composition listed in Tables S1 and S2. (D) cPLA_2_-membrane association and dissociation processes.

We simulated cPLA_2_ with and without bound Ca^2+^ at the C2 domain. The C2 domain is composed of an 8-stranded anti-parallel β-sandwich. The changes in the RMSF of the several secondary structures at the C2 domain (Figure 2A and 2B) were measured. RMSF quantifies the average deviation of the cPLA_2_ Cα from its mean position during the simulations. Our simulations agree with data for Ca^2+^ binding at the C2 domain, which rigidifies, stabilizes, and activates cPLA_2_. The association of the C2 domain with the phospholipid membrane requires Ca^2+^.^23^

**Figure 2.**
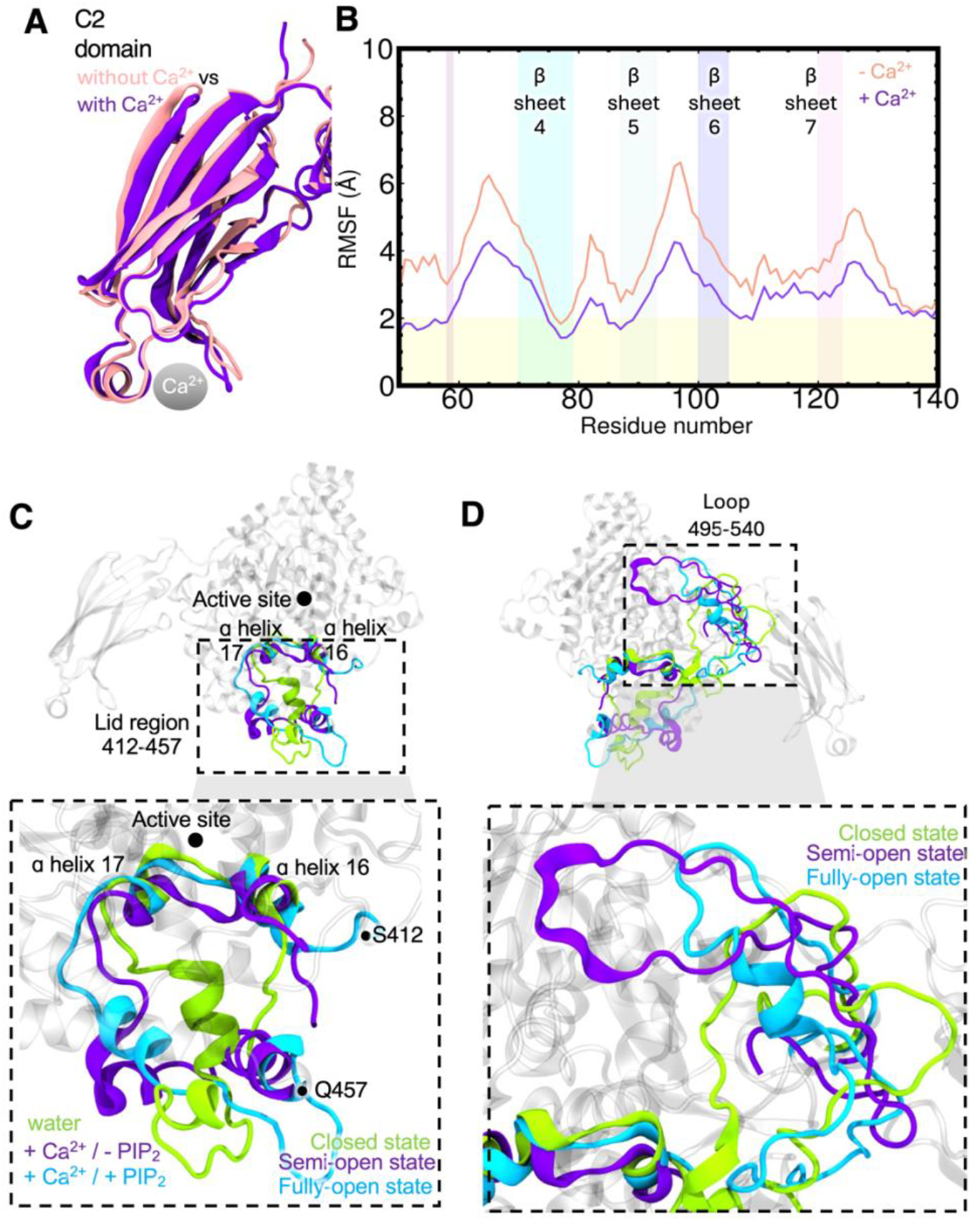
Conformational dynamics of C2 domain, lid region, and loop 495-540 cPLA_2._ (A) Superimposed C2 domain of system without and with Ca^2+^; (B) RMSF of β sheets 4, 5, 6, and 7 (right). RMSF for the Cα atoms were measured from the n=3 replicates of 500 ns all-atom MD simulations. Top clustered conformation of aqueous cPLA_2_ (green); cPLA_2_ / + Ca^2+^ / - PIP_2_ (purple); and cPLA_2_ / + Ca^2+^ / + PIP_2_ (cyan). Closed, semi-open, and fully-open states for the lid region and loop 495-540 are shown. (C) Superimposed lid region residues 412-457; and (D) Superimposed loop region residues 495-540.

Ca^2+^ binding neutralizes the negative electrostatic potential at the loop and β-sheet residues. The hydrophobic interactions between the membrane and the C2 domain allow the association of the now neutral C2 environment due to the lower free energy cost of inserting charged amino acids into the membrane.^24^ Furthermore, increased Ca^2+^ concentration signals the translocation of cPLA_2_ from the cytosol to the plasma membrane, Golgi, and ER. We further investigated how the catalytic domain also recognizes the membrane surface.^25^

Our other goal was to understand how the well-known allosteric regulator, phosphatidylinositol (4,5)-bisphosphate (PIP_2_), affects the activation process. PIP_2_ is predominantly concentrated in the plasma membrane, which is a critical regulator of cell signaling. However, a small yet measurable fraction (∼1%) exists in the endoplasmic reticulum (ER), not as a signaling hub but as an intermediate in lipid metabolism. The ER primarily functions as a lipid-processing center, supplying PI for PIP_2_ biosynthesis at the plasma membrane, thereby maintaining the dynamic equilibrium essential for cellular signaling fidelity. In our MD simulations, PIP_2_ continuously diffuses laterally in the membrane. This implies that PIP_2_ remained resistant to extraction in the catalytic site of cPLA_2_, likely due to its highly negative polarity and the intrinsic timescale of lipid binding exceeding the 500 ns simulation timescale. This suggests that while the portion of the PIP_2_ in the membrane may associate with the PIP_2_ allosteric sites, this was not observed but would requires extended simulation times to overcome the slower, collective lipid reorganization processes essential for binding.

The lid region samples multiple conformations that can interconvert between the closed, semi-open, and open conformations. HDX-MS was used to probe how the different parts of cPLA_2_, including the lid region, interact with the membrane.^19,20^ Through a series of HDX-MS experiments, the lid region was found to be a flexible regulated structure with a particular orientation to the membrane.^19^ After further structural and conformational dynamics analysis of different cPLA_2_ systems, we found that the lid region and loop 495-540 showed high fluctuations throughout the simulations (Figures 2C and 2D). It should be noted that a part of the lid region (433-459) is not solved in the X-ray structure. However, after running multiple replicates of 500 ns equilibrium MD, the lid region stabilizes as measured by the protein backbone RMSD (Figure S2).

After further investigation, we checked the lid region and noted several key differences at the C-terminal end (435-457) (Figure 3). First, we overlapped the highest clustered conformation of aqueous cPLA_2_ and the fully activated cPLA_2_ (+Ca^2+^ / +PIP_2_) (Figure 3A). The lid region (412-457) is composed of two α helices (α helix 16 and α helix 17) and a loop (435-457). It is expected that the helices will be more stable than the loop. The fully activated form cPLA_2_ (+Ca^2+^ / +PIP_2_) has the most stable loop 435-457. Interestingly, this loop 435-457 fluctuates up to two-fold more in an aqueous environment (Figure 3B). Second, the membrane association stabilized these fluctuations (Figure 3A and 3B). The fully activated form of cPLA_2_ that has bound Ca^2+^ and with accessible PIP_2_ in the membrane was found to have the least RMSF, while the aqueous cPLA_2_ has the highest RMSF in the 435-457 region. Third, we noted several other conformational states of the lid region (Figure 3C, 3D, 3E, 3F) that we classified as closed, semi-open, and fully open states. Aqueous cPLA_2_ is in a closed state, while the fully activated form of cPLA_2_ that has bound Ca^2+^ and accessible PIP_2_ is in a fully open state.

**Figure 3.**
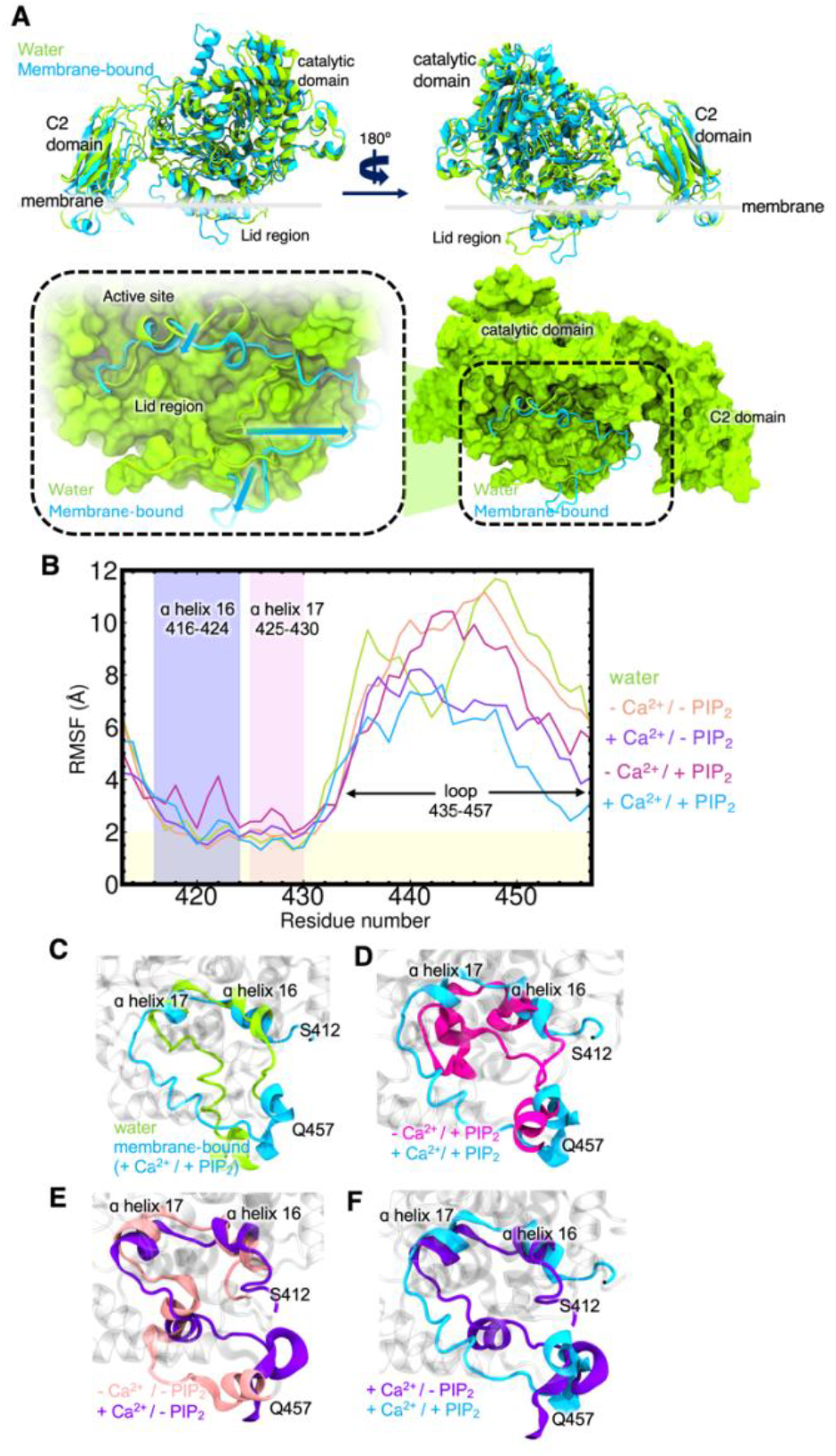
Lid region dynamics. (A) Lid region conformational dynamics after membrane association; (B) RMSF of α helix 16, α helix 17, and loop 435-457. RMSF for the Cα atoms were measured from the combined n=3 replicates of 500 ns all-atom MD simulations. The superimposed top clustered conformer of each system (C) water vs. +Ca^2+^ / +PIP_2_; (D) -Ca^2+^ / +PIP_2_ vs +Ca^2+^ / +PIP_2_ ; (E) -Ca^2+^ / -PIP_2_ vs. +Ca^2+^ / -PIP_2_ ; and (F) +Ca^2+^ / -PIP_2_ vs +Ca^2+^ / +PIP_2_. RMSF for the Cα atoms were measured from the n=3 replicates of 500 ns all-atom MD simulations. All non-aqueous systems are membrane-associated.

cPLA_2_ features a deep, channel-like active site shielded by a lid region. Additionally, the lid and the areas surrounding active site residues S228 and D549 are enriched with aromatic amino acids, which favorably recognize and bind the double bonds in the 20:4 arachidonic acid tail via π-π stacking. The X-ray structure (PDB ID 1CJY) is presumed to represent the “closed” form of cPLA_2_. Although it is partially activated with Ca^2+^ bound in the C2 domain, the fact that a fatty acyl chain lacks direct access to the active site confirms that cPLA_2_ also needs further conformational changes. Upon membrane association, it appears that the active site will become more accessible, which we confirmed in our MD simulation data.

Using principal component analyses (PCA), we detected significant movement of the loop 495-540 (Figure 4A and Figure 4B). The goal of PCA in molecular simulations of protein dynamics is to identify the most dominant and collective motions of a protein by reducing the dimensionality of the trajectory data. Here, the Cα RMSD was used in the PC calculations, and the two most dominant PCs, PC1 and PC2, were projected onto the cPLA_2_ structure. Interestingly, the free energy minimum of aqueous cPLA_2_ is adjacent to that of fully activated cPLA_2_ (+Ca^2+^/ +PIP_2_), suggesting that the lid region can readily transition between open and closed states. Additionally, the simulation movie revealed a lateral displacement of the loop spanning residues 495–540 (as captured by PC1). This movement appears to facilitate the exposure of the allosteric lysine residues K488, K541, K543, and K544 following membrane association and activation.

**Figure 4.**
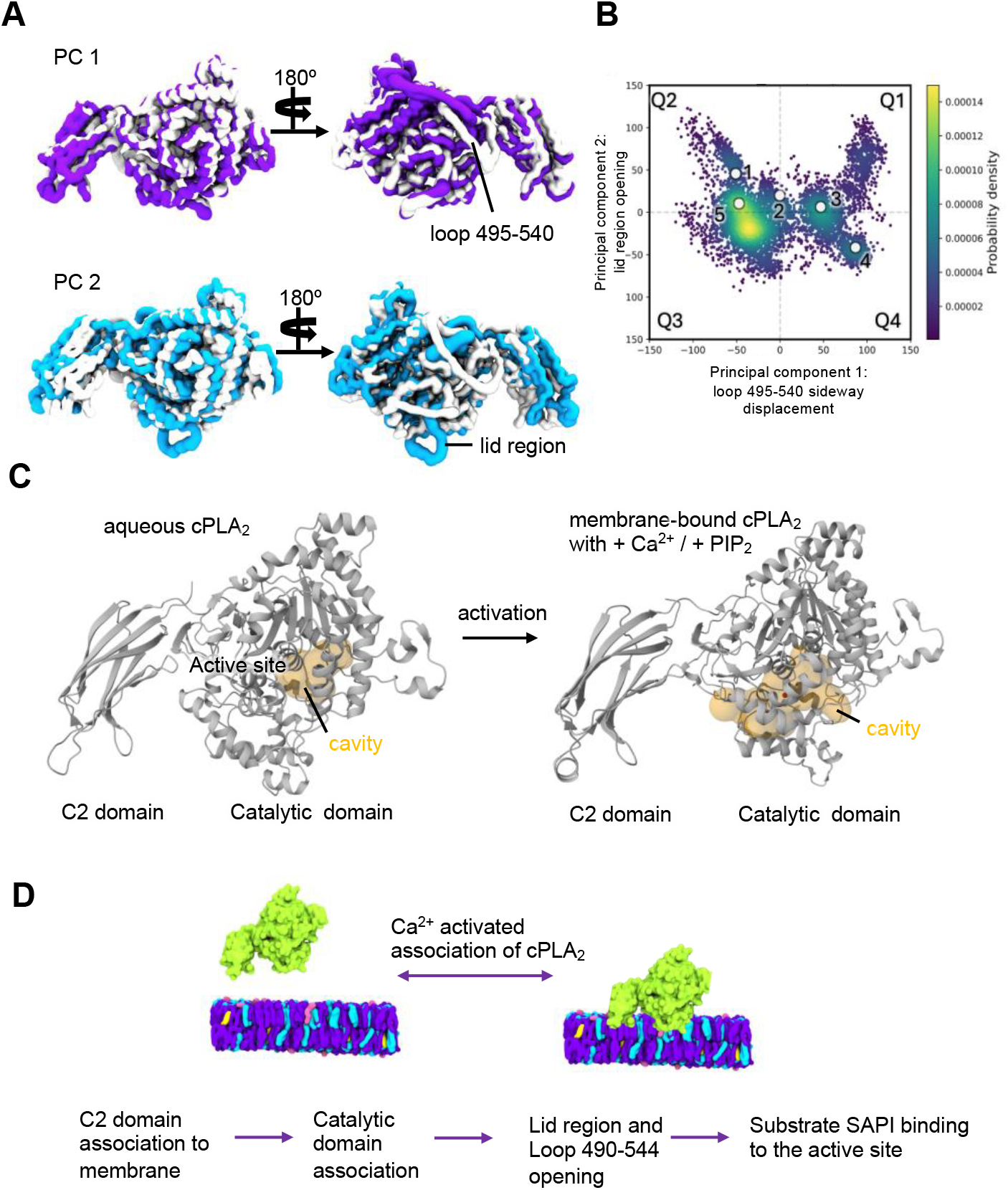
Mechanistic dissection of cPLA_2_ activation. (A) Projection of Principal components 1 and 2 to the protein structure. The X-ray structure (gray) was superimposed in the last frame of PCs 1 and 2. (B) Probability density of all the PCs from system **1** aqueous cPLA_2_ ; **2** cPLA_2_ / - Ca^2+^ / - PIP_2_; **3** cPLA_2_ / + Ca^2+^ / - PIP_2_; **4** cPLA_2_ / - Ca^2+^ / + PIP_2_; **5** cPLA_2_ / + Ca^2+^ / + PIP_2_; (C) Evolution of phospholipid substrate tunnel inside the catalytic domain active site. Cavity volume was calculated and visualized with the CAVER3.0 web server using a value of 0.1 for minimum probe radius. (D) cPLA_2_ membrane association with Ca^2+^. Proposed mechanism of active site opening: C2 domain association with the membrane; Catalytic domain associates; Lid region and loop 490-544 transition from closed-state to open-state; and active site opened to accommodate SAPI substrate.

In human cPLA_2_, the lid region is composed of 23.9% negatively charged residues (D and E), 9% positively charged residues (R and K), and 28% of hydrophobic residues (Figure S1). Sequence alignment of human, monkey, rabbit, and mouse cPLA_2_ shows that this region shares 85–98% identity across these species, with chicken being an exception at 70% identity. Notably, the basic residues are 67% conserved, and the acidic residues are 81.2% conserved, highlighting the strong evolutionary conservation of the lid region. For RAW264.7 macrophage membranes, which carry a net negative charge, the 23.9% acidic composition of the lid likely facilitates its repulsion and the spontaneous exposure of the active site. We propose that the lid’s amphipathic nature promotes membrane association and enhances substrate recognition and binding, particularly in environments where the membrane patch exhibits variable net charge.

The transition of cPLA_2_ from a partially activated form to its fully activated form is also characterized by a three-fold increase of the active site cavity volume compared to the aqueous cPLA_2_ (closed state) around the catalytic dyad residues, S228 and D549 (Figure 4C). We used the web server tool CAVER 3.0 to detect the cavity size around the active site.^26^ The allosteric site residue K544 also gets exposed to the cavity after membrane association. K544 exposure is not observed during simulations of aqueous cPLA_2_. Furthermore, with CAVER cavity detection analyses, we observed that the active site became more available and thus should allow easier access for phospholipid substrates.

We propose a mechanistic event following the membrane association (Figure 4D). Based on our all-atom MD simulations data, we deduced that the active site opening started with (step 1) C2 domain association with the membrane; (step 2) catalytic domain association; (step 3) lid region and loop 490-544 transitioned from a closed-state to an open-state; and (step 4) the active site opens to accommodate the SAPI substrate.

The lid region opening of cPLA_2_ is an allosteric hub in a population-shift allostery model. This model posits that the protein’s free energy landscape comprises an ensemble of conformations. In contrast to its behavior in aqueous environments, membrane-associated cPLA_2_ exhibits a more open lid region, with a small probability of transitioning to a closed state. The additions of Ca^2+^ and PIP_2_ at the C2 domain and membrane strengthen the lid opening effect.

Although our simulations are limited and may not capture the phospholipid substrate SAPI or PIP_2_ binding, we surmised that we sampled enough events to characterize the lid region opening and loop conformational plasticity after membrane association. Not only did MD simulations help refine the loop structures, but they also suggested the deeper biophysical basis of its role in cPLA_2_ membrane-association and activation. It should be noted that simulating protein loop dynamics remains one of the challenges in membrane protein modeling.

In summary, we show how the membrane-associated cPLA_2_ transitions from closed to open by opening its lid region. Ca^2+^ allows the stabilization of the C2 domain after cPLA_2_ associated with the membrane that verifies published experimental data. Furthermore, we see how the movement of loop 495-540 preceding the allosteric site activation can be crucial for PIP_2_ recognition. Lastly, this more profound understanding of the cPLA_2_-membrane environment can help illuminate the foundation for its mechanism of interaction with the membrane and the discovery of better therapeutics for inflammatory diseases based on active and/or allosteric site inhibition of cPLA_2_.

## Abbreviations

AA: arachidonic acid
cPLA_2_: Group IVA (GIVA) cytosolic phospholipase A_2_
ER: Endoplasmic reticulum
HDX-MS: Hydrogen/Deuterium exchange-mass spectrometry
MD: molecular dynamics
PAPC: 1-palmitoyl-2-arachidonoyl-*sn*-glycero-3-phosphocholine
PC: phosphatidylcholine
PCA: principal component analysis
PLA_2_: Phospholipase A_2_
PI: phosphatidylinositol
PIP_2_: phosphatidylinositol (4,5)-bisphosphate
PS: phosphatidylserine
RMSF: Root-mean-square fluctuation
RMSD: Root-mean-square deviation
SAPI: 1-stearoyl-2-arachidonoyl phosphatidylinositol

## Supporting Information

Computational details, Tables S1-S3, Figures S1–S3, and Movie S1.

## Notes

The authors declare no competing financial interest.

## Data availability

Trajectories, structures, simulation scripts, analysis scripts, and data files are uploaded to our group’s website and can be accessed with this link https://amarolab.ucsd.edu/data.php.

## Author contributions

M.K.E.B., E.A.D., and R.E.A. designed the project. M.K.E.B. performed the modeling, simulations, and analyzed the data. E.A.D. and R.E.A. supervised the modeling and MD simulations. M.K.E.B. wrote the initial draft of the paper. E.A.D. and R.E.A. secured the funding and resources for the project. All authors contributed to the writing and editing of the manuscript.

## Acknowledgments

We thank Dr. Nicolas Frederic-Lipp, Dr. Abigail Dommer, Prof. Itay Budin, Prof. J. Andrew McCammon, Dr. Varnavas Mouchlis, Dr. Daiki Hayashi, Dr. Gosia Murawska, Dr. Carla Calvó-Tusell, Dr. Lorenzo Casalino, Dr. Mohamed Shehata, Xandra Nuqui, Clare Morris, and Nicholas Wauer, for helpful discussions and advice on phospholipase A_2_, simulations, and analysis. Supercomputing resources were provided to R.E.A. by XSEDE (NSF TG-CHE060073) and ACCESS (NSF TG-CHE060063) allocations. The study was supported by National Institutes of Health grants NCI P01-CA234228 (R.E.A.) and R35 GM139641 (E.A.D.).

## Graphical TOC

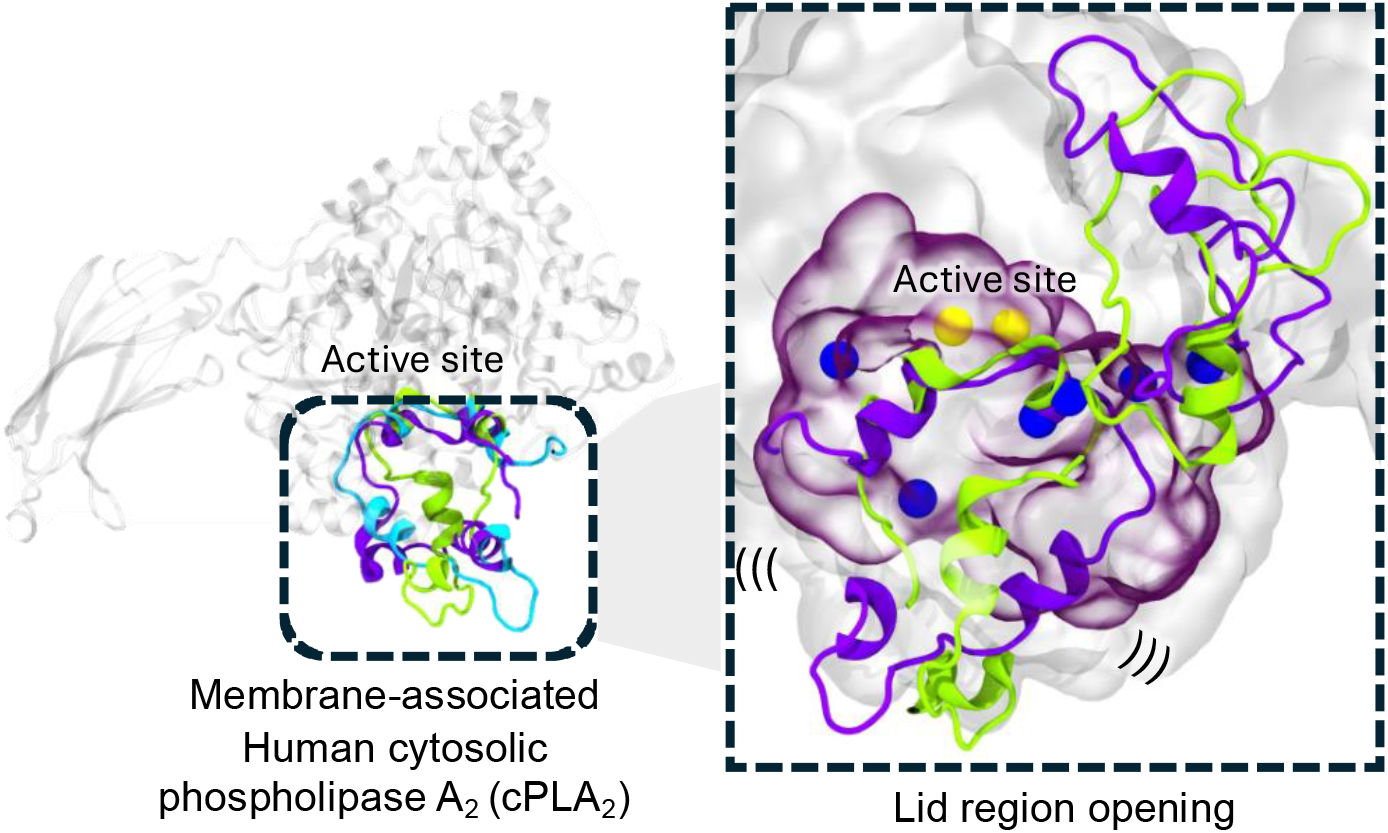

