## Supplementary Information for "Conformational dynamics and activation of membrane-associated human Group IVA cytosolic phospholipase A_2_ (cPLA_2_)"

### Table of Contents

**Sections** **Page No.**

**Computational methods** **2**

- Human Group IVA $cPLA_2$ modeling
- RAW264.7 Macrophage Endoplasmic Reticulum Membranes Modeling
- All-atom molecular dynamics simulations
- Trajectory analyses

**Tables** **3-4**

- Number of lipids present in lipid bilayer membrane without $PIP_2$
- Number of lipids present in lipid bilayer membrane with 1% $PIP_2$
- cPLA2 secondary structure

**Figures** **5-7**

- Lid region sequence conservation and conformational dynamics.
- Multiple sequence alignment of different $cPLA_2$ loop 495-545 from various species.
- Lid region RMSD.
- Loop 495-540 region RMSF.
- Modeled $cPLA_2$-SAPI enzyme-substrate complex.

**Movies** **7**

- Loop and lid region movement of aqueous $cPLA_2$.
- Loop and lid region movement of membrane-associated $cPLA_2$.

**References** **7**

### Computational Methods

#### Human Group IVA $cPLA_2$ modeling

The available human $cPLA_2$ X-ray structure (PDB ID: 1CJY) was used. It has both the N-terminal calcium-dependent lipid-binding C2 and catalytic domains.[1] Non-terminal loops missing residues were modeled in the CHARMM-GUI web server PDB Reader. GIVA $cPLA_2$ chain A has three missing non-terminal loops that were included in the model: residues 406-414, 431-457, and 498-537. CHARMM-GUI employed the GalaxyFill algorithm in modeling the non-terminal missing residues.[2,3] The calcium bound in the $cPLA_2$ crystal structure was also modeled and parameterized. Amino acid residues were assigned their standard protonation state at pH 7 using PROPKA.[4] A 0.15 M $Na^+$ and $Cl^-$ ions at a concentration of 0.15 M were added to neutralize the system by using the Monte-Carlo ion placing method in the CHARMM-GUI web server.

#### RAW264.7 Macrophage Endoplasmic Reticulum Membranes Modeling

The ER lipid bilayer model was generated using the CHARMM-GUI Membrane Builder.[5] The RAW264.7 macrophage ER membrane composition was based on the published subcellular lipidomics data of the Dennis group.[6] The complete ratio of all lipids used in our membrane is reported in **Table S1 and S2** for a membrane without and with 1% $PIP_2$, respectively. In CHARMM-GUI Membrane Builder, a water thickness of 50 Å covering the whole protein on top of the membrane was used. PPM 2.0 was also used to ensure that the protein is correctly aligned with respect to the membrane based on the Orientation of Protein in a Membrane web server database. The length of X and Y dimensions was set to 150 Å. The box size was 156.1 x 156.1 x 213.3 $Å^3$. A 0.15 M $Na^+$ and $Cl^-$ ions were added (551 $Na^+$ and 359 $Cl^-$) to neutralize the system. There are 400 lipids in both the upper leaflet and lower leaflet of the ER membrane.

#### All-atom molecular dynamics simulations

GPU-enabled NAMD 3.0 was used for all-atom MD simulations of the aqueous and membrane-associated $cPLA_2$.[7] Each replicate was carried out for 500 ns production runs with three replicates each in National Center for Supercomputing Applications (NCSA) Delta. NVIDIA A100 GPU nodes were used. The CHARMM36 force field was used for the protein and lipids.[8] The TIP3P water model was used for solvent.[9] The system was minimized for $5x10^6$ steps (with 1fs/step and for a total of 5 ns) and then followed by an equilibration of $5x10^6$. Velocities were set randomly from a Boltzmann distribution and differed in each replicate. During equilibration, constant temperature was set to 310 K with Langevin dynamics. A 12 Å cutoff was used to calculate electrostatic interactions. For the CHARMM36 force field the scaled 1-4 was used for the nonbonded parameters. Production was run with NPT ensemble by using 1.01325 bar pressure and 310 K temperature and using 2 fs/step timestep. Each frame was outputted every 1 ns.

#### Trajectory Analyses

The first 30 ns of production runs were discarded to make sure that the systems are well-equilibrated. VMD was used to visualize trajectories and render all the protein and membrane structures.[10] Each $cPLA_2$ trajectory was aligned to its frame by Cα atoms to remove rotational and translational degrees of freedom.

Root mean square deviation (RMSD) and root-mean-square fluctuation (RMSF) calculations of protein backbone atoms were performed in pytraj. Plots were rendered using python's matplotlib library.[11]

Principal component analysis of the $cPLA_2$ Cα atoms were calculated using cpptraj.[12] The covariance matrix was obtained from the atomic fluctuations of the Cα's and diagonalized to yield eigenvectors and eigenvalues. These represent the direction and magnitude of motions, respectively. The two highest eigenvalues were reported as PC 1 and PC 2 and were projected back to Cartesian coordinate. k-means clustering method was applied to obtain the top visited structural conformation. Backbone RMSD of protein residues were used as distance metric. Sieve value was set to 10. Topmost clustered structures were compared. Using the topmost clustered structures, we used CAVER3.0 web server to calculate and identify the volume of the active site cavity.[13]

**Table S1**. Number of lipids present in lipid bilayer membrane without $PIP_2$.

| Lipids | Count | |
|---|---|---|
| | **Upper leaflet** | **Lower leaflet** |
| Cholesterol | 20 | 20 |
| DPPC | 8 | 8 |
| POPC | 24 | 24 |
| PLPC | 8 | 8 |
| SLPC | 48 | 48 |
| POPE | 24 | 24 |
| SOPE | 28 | 28 |
| DOPE | 44 | 44 |
| POPS | 40 | 40 |
| SAPI | 44 | 44 |
| PSM | 48 | 48 |
| BSM | 4 | 4 |
| Cer6 | 8 | 8 |
| CER6X | 12 | 12 |
| CER160 | 4 | 4 |
| PLA20(PE) | 36 | 36 |
| **Total number of lipids** | 400 | 400 |

**Table S2**. Number of lipids present in lipid bilayer membrane with 1% $PIP_2$.

| Lipids | Count | |
|---|---|---|
| | **Upper leaflet** | **Lower leaflet** |
| Cholesterol | 20 | 20 |
| DPPC | 8 | 8 |
| POPC | 24 | 24 |
| PLPC | 8 | 8 |
| SLPC | 48 | 48 |
| POPE | 24 | 24 |
| SOPE | 28 | 28 |
| DOPE | 44 | 44 |
| POPS | 40 | 40 |
| SAPI | 40 | 40 |
| SAPI24 | 4 | 4 |
| PSM | 48 | 48 |
| BSM | 4 | 4 |
| Cer6 | 8 | 8 |
| CER6X | 12 | 12 |
| CER160 | 4 | 4 |
| PLA20(PE) | 36 | 36 |
| **Total number of lipids** | 400 | 400 |

**Table S3**. $cPLA_2$ secondary structure.

| Residue numbers | Description |
|---|---|
| 18 - 28 | Beta sheet 1 |
| 33 - 40 | Helix 1 |
| 44 – 50 | Beta sheet 2 |
| 58 - 59 | Beta sheet 3 |
| 70 - 79 | Beta sheet 4 |
| 87 – 93 | Beta sheet 5 |
| 100 – 105 | Beta sheet 6 |
| 120 – 124 | Beta sheet 7 |
| 128 – 134 | Beta sheet 8 |
| 151 – 174 | Helix 2 |
| 179 – 182 | Helix 3 |
| 190 - 194 | Beta sheet 9 |
| 197 – 215 | Helix 4 |
| 221 – 226 | Beta sheet 10 |
| 227 – 240 | Helix 6 |
| 262 – 267 | Helix 7 |
| 268 – 285 | Helix 8 |
| 290 – 305 | Helix 9 |
| 306 – 309 | Helix 10 |
| 312 – 319 | Helix 11 |
| 326 – 333 | Beta sheet 11 |
| 340 – 344 | Helix 12 |
| 345 – 349 | Beta sheet 12 |
| 354 – 355 | Beta sheet 13 |
| 362 – 363 | Beta sheet 14 |
| 364 – 368 | Helix 13 |
| 372 – 373 | Beta sheet 15 |
| 376 – 377 | Beta sheet 16 |
| 385 - 394 | Helix 14 |
| 395 – 400 | Helix 15 |
| 416 – 424 | Helix 16 |
| 425 – 430 | Helix 17 |
| 462 – 477 | Helix 18 |
| 488 – 490 | Beta sheet 17 |
| 544 – 548 | Beta sheet 18 |
| 549 – 553 | Helix 19 |
| 557 – 562 | Helix 20 |
| 570 – 575 | Beta sheet 19 |
| 587 – 600 | Helix 22 |
| 610 – 616 | Helix 23 |
| 620 – 623 | Beta sheet 20 |
| 636 – 641 | Beta sheet 21 |
| 645 – 648 | Helix 24 |
| 650 – 652 | Beta sheet 22 |
| 655 – 656 | Beta sheet 23 |
| 659 – 667 | Helix 25 |
| 686 – 703 | Helix 26 |
| 704 – 719 | Helix 27 |

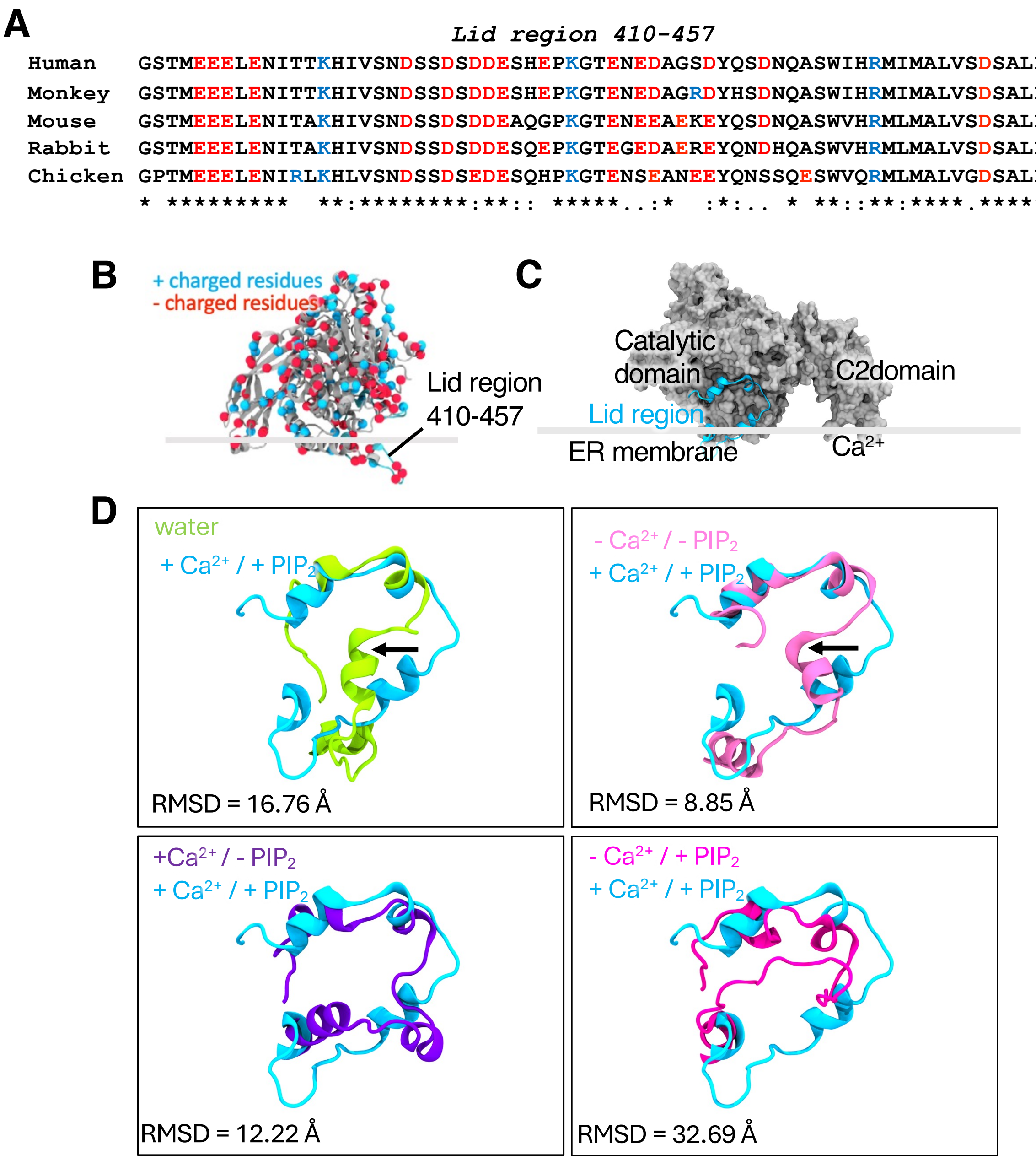


**Figure S1**. Lid region sequence conservation and conformational dynamics. (A) Multiple sequence alignment of different $cPLA_2$ from various species. Human *Homo sapiens* (Uniprot ID P47712); Ma's night monkey *Aotus nancymaae* (Uniprot ID A0A2K5ETQ0); Mouse *Mus musculus* (Uniprot ID P47713); Rabbit *Oryctolagus cuniculus* (Uniprot ID Q9TT38); Chicken *Gallus gallus* (Uniprot ID P49147). (B) Distribution of positive (blue) and negative (red) charges amino acid in $cPLA_2$. (C) Lid region orientation towards the ER membrane. (D) Superimposition of the topmost clustered conformation of $cPLA_2$ with +/- Ca2+ and +/- PIP2. The RMSD values were calculated on the protein backbone atoms with the $cPLA_2$ + $Ca^{2+}$ and + $PIP_2$ was used as reference.

loop 495-544

```
Human    TREGRAGKVHNFMLGLNLNTSYPLSPLSDFATQDSFDDDELDAAVADPDEFERIYEPLDVKSKK  544
Monkey   TREGRAGKVHNFMLGLNLNTSYPLSPLSDFATQDSFDDDELDAAVADPDEFERIYEPLDVKSKK  544
Mouse    TREGRAGKVHNFMLGLNLNTSYPLSPLRDFSSQDSFD-DELDAAVADPDEFERIYEPLDVKSKK  543
Rabbit   TREGRAGKVHNFMLGLNLNTSYPLSPLRDFTQESFDD-DELDAAVADPDEFERIYEPLDVKSKK  543
Chicken  TREGRAGKVHNFMLGLNLNSCYPLSPLADLLTQESVEEDELDAAVADPDEFERIYEPLDVKSKK  544
         ******************:.****** *:  :.  : *************************
```

**Figure S2**. Multiple sequence alignment of different cPLA$_2$ loop 495-545 from various species. Human *Homo sapiens* (Uniprot ID P47712); Ma's night monkey *Aotus nancymaae* (Uniprot ID A0A2K5ETQ0); Mouse *Mus musculus* (Uniprot ID P47713); Rabbit *Oryctolagus cuniculus* (Uniprot ID Q9TT38); Chicken *Gallus gallus* (Uniprot ID P49147).

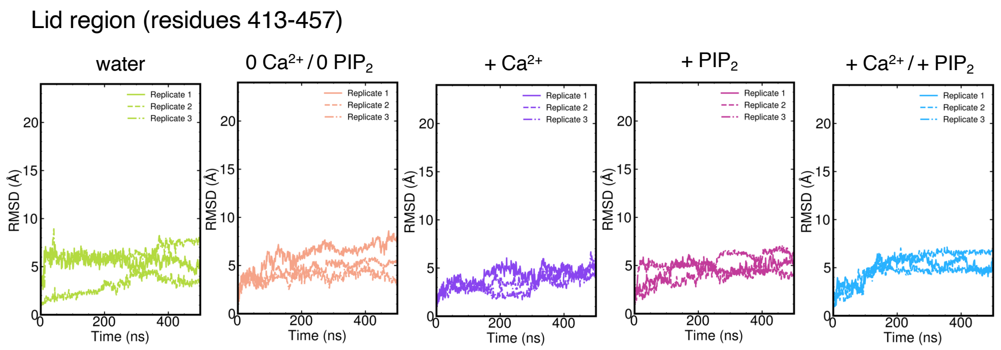


**Figure S3**. Lid region RMSD.

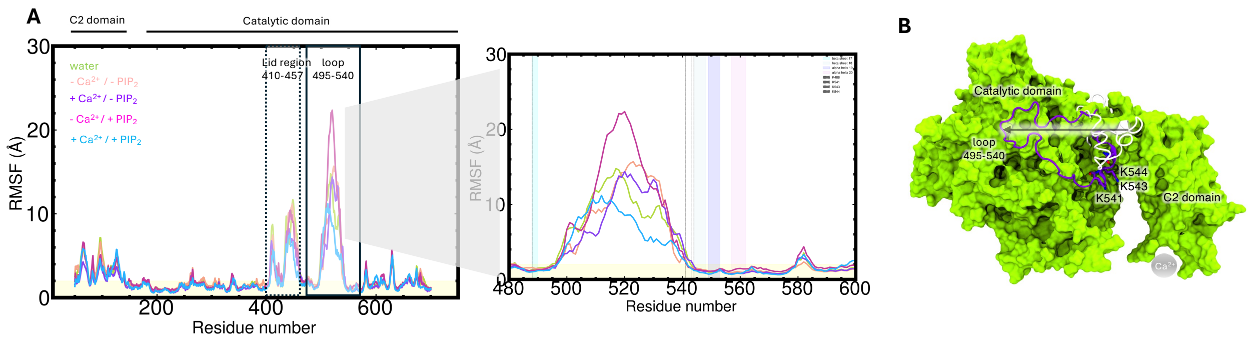


**Figure S4**. Loop 495-540 region RMSF. (A) RMSF plot of all systems simulated; and (B) Sideway displacement of loop 495-540.

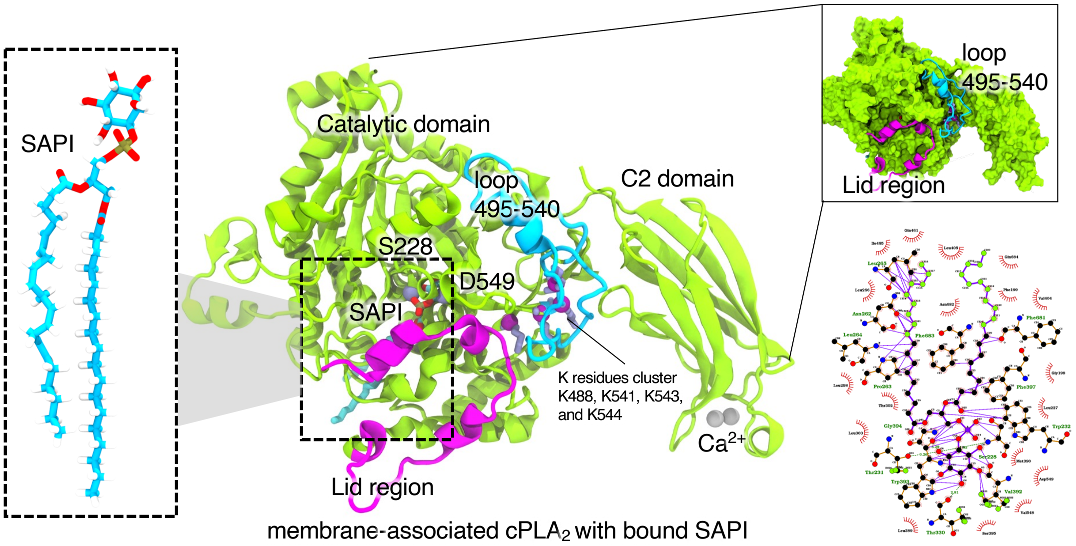


**Figure S5**. Modeled $cPLA_2$-SAPI enzyme-substrate complex.

**Movie S1**. Loop and lid region movement of aqueous $cPLA_2$.
**Movie S2**. Loop and lid region movement of membrane-associated $cPLA_2$.
